# Exogenous melatonin mitigates drought stress by promoting the living cortical area of roots in cotton

**DOI:** 10.1101/2025.02.28.640717

**Authors:** Haitao Zhang, Congcong Guo, Hongchun Sun, Lingxiao Zhu, Ke Zhang, Yongjiang Zhang, Zhanbiao Wang, Cundong Li, Liantao Liu

**Author notes:** **Correspondence:** (C.L.); (L.L.). Haitao Zhang and Congcong Guo contributed equally to this work.

## Abstract

The root cortex plays a critical role in water absorption, influencing root function and metabolic activity. However, the effects of melatonin on root cortical cells and its role in drought resistance remain unclear. This study examines the impact of exogenous melatonin on the root cortex of cotton under drought stress, focusing on its relationship with water uptake and drought resilience. Cotton plants were treated with 8% PEG6000 for drought stress and 100 µM melatonin by foliar application. The results demonstrated that melatonin application significantly increased the living cortical area of roots under drought stress, with notable increases of 26.34% and 59.00% at distances of 1 cm and 13 cm from the root tip, respectively. Melatonin also enhanced root cortical thickness, promoted osmotic regulation, increased respiratory enzyme activity, and improved water and nutrient uptake. Furthermore, melatonin application promoted both root and shoot growth. Notably, the living cortical area positively correlated with osmotic substance accumulation, root respiration, and absorption capacity under drought conditions. In conclusion, exogenous melatonin alleviates drought-induced damage by enhancing the living cortical area, optimizing osmotic regulation, and improving root absorption, thus promoting cotton growth and drought resistance. These findings highlight melatonin as a promising regulator for enhancing drought resistance.

**Highlight:** Exogenous melatonin increases root cortical integrity of cotton roots under drought stress, improves the absorption of water and nutrients, promotes the growth of cotton, and enhances its drought resistance.

## Introduction

Cotton is an important economic crop, fiber crop, and oilseed crop (Dong et al., 2008). Drought has become a key limiting factor affecting cotton production, severely impacting its growth, development, and yield (Chen & Dong, 2016). With the intensification of global climate change, the frequency and severity of drought events are bound to increase, further exacerbating water stress in cotton production, especially in rain-fed agricultural areas (Ma et al., 2018). Therefore, enhancing the drought resistance of cotton is essential.

The structure and anatomical features of plant roots exhibit plasticity, which is crucial for adapting to drought stress. Drought stress can directly affect root growth. Research by Yang et al. (2023) shows that drought stress significantly reduces total root length, total total root surface area, total root volume, average root diameter, and lateral root branching density in cotton. The root cortical tissue, particularly in regions with strong absorptive functions, constitutes a significant portion of the root system and serves as a vital site for the synthesis, storage, and secretion of nutrients and growth-regulating substances (Lynch, 2015). However, the cortex can consume over 50% of photosynthetic products to sustain survival, making it the primary region of the root system where photosynthetic resources are expended. Therefore, changes in the root cortex under drought stress are critical for the adaptation of plant roots to drought conditions (Lambers et al., 2006).

The anatomical structure of the cortex is closely related to root metabolic consumption. Under soil stress conditions, the plant’s anatomical structure changes, adjusting the proportion of respiring and non-respiring root tissue to adapt to or alleviate adverse conditions (Schneider & Lynch, 2018). In short, drought stress diminishes plant productivity, disrupts the integrity of root cortical cells, reduces cellular metabolic activity, thereby inhibiting root respiration and decreasing root metabolic consumption (Chimungu et al., 2014). This indicates that the amount of living cells in the root cortex under drought stress is closely related to root respiratory metabolism. Structure influences function; the root cortex serves as a lateral transport pathway for water and nutrients into the vascular cylinder. A reduction in the living cortical area of the roots will lead to decreased absorption capacity, thereby affecting water and nutrient uptake (Zhan & Lynch, 2015). Hu et al. (2014) found that as the living cortical area of maize decreases, the radial transport of phosphates, calcium, and sulfates in maize roots also decreases. Therefore, an early reduction in the root cortical area can significantly regulate the source-sink relationship between the roots and aerial parts, resulting in stunted growth of the aerial portion, reduced photosynthetic capacity, and overall detriment to the plant.

Drought stress severely limits cotton production, and enhancing drought resistance is crucial for yield stability (Lobell et al., 2014). Exogenous plant growth regulators, such as melatonin, have shown promise in improving crop drought resistance (Guo et al., 2023). Melatonin is widely present in plants and acts as a plant growth regulator that enhances resistance to various biotic and abiotic stresses (Sun et al., 2021; Wang et al., 2022). Applying melatonin can promote root elongation in cotton and increase the number of lateral roots, thereby expanding the total root surface area and volume, resulting in a richer root system that fosters plant growth (Zhu et al., 2023). Under drought stress, the application of exogenous melatonin significantly increases the accumulation of osmotic regulatory substances in the roots of hickory, providing protective effects and ensuring that the roots can maintain normal physiological functions in drought conditions (Wang et al., 2019). Thus, the root system is considered an important tissue through which melatonin enhances plant stress resistance. The root cortex affects root activity and is a crucial site for reflecting the absorption function of the roots. Its functionality and physiological metabolism levels largely represent the overall functioning and metabolic status of the root system. Therefore, it is hypothesized that the regulatory effects of melatonin on the roots primarily target the cortical tissue. Previous studies have found that the application of melatonin under stress conditions significantly impacts the anatomical structure of plant roots. Research indicates that under salt stress, the cortical cells of cotton are the first to experience damage, exhibiting severe membrane rupture; however, melatonin can help maintain the structural integrity of the cortical cells, effectively alleviating damage caused by salt stress (Duan et al., 2022). Under drought stress, the application of melatonin significantly increases the size of the root epidermis, pith, cortex, secondary xylem, and vascular bundles in *Eugenia uniflora*, restoring them to their natural homeostasis (David et al., 2023). Hence, it is speculated that the exogenous application of melatonin would have a positive effect on the root cortex of cotton under drought stress.

The application of exogenous melatonin has been shown to mitigate the adverse effects of drought stress on plants and to influence the anatomical structure of the root system. However, few studies have investigated whether melatonin affects root metabolic processes and water and nutrient absorption by influencing cortical cells. Therefore, this study hypothesizes that under drought stress, exogenous melatonin can increase the living cortical area of roots, prolong the functional integrity of the root cortex, promote cotton growth, and enhance drought resistance. The main objectives of this research are: (1) to elucidate the characteristics, patterns, and interrelationships of melatonin’s effects on the area of living cells in the root cortex, osmotic regulatory substance content, and respiration rate in cotton roots under drought stress; (2) to reveal the impact of the area of living cortical cells on root growth and development in cotton under drought stress, along with their interrelationships; (3) to explore how the area of living cortical cells in cotton roots under drought stress influences drought resistance in aerial parts and the relationships involved.

## Materials and Methods

### Experimental design

The experiment was conducted from January to October 2024 in the artificial climate chamber at Hebei Agricultural University in Baoding City, Hebei Province (38°85′N, 115°30′E). The cotton variety used in the study was “Lumian 532.” Uniform and fully developed cotton seeds were selected, disinfected by soaking in 75% alcohol for 10 minutes and then rinsed with distilled water 5-8 times to remove all residual alcohol. After rinsing, the seeds were placed in distilled water and kept in a dark constant-temperature incubator at 25 °C for 8 hours. The seeds were then spread out on a tray covered with a damp towel to germinate for 24 hours. Seeds with radicles approximately 1 cm long were transplanted onto a foam board with holes for hydroponic growth. Once the cotyledons fully expanded, seedlings with intact cotyledons and consistent growth were transplanted into hydroponic boxes (43×30×20 cm). Before use, the boxes were thoroughly disinfected with 50% carbendazim and then rinsed 2-3 times with clean water. In the hydroponic boxes, Hoagland nutrient solution was used, which was replaced every 3 days, and aeration was provided twice daily for 1 hour each time. The artificial climate chamber was set at 25/22 °C (day/night), 600 μmol/m²/s light intensity, 14 hours photoperiod, and 60±5% relative humidity.

When the seedlings developed three true leaves, uniform plants were selected for water and melatonin (MT) foliar spray treatments: CK (Well water, 0% PEG 6000), DS (Drought stress, 8% PEG 6000), CK+MT (Well water and 100 μmol/L MT), and DS+MT (Drought stress and 100 μmol/L MT). During the treatment period, the CK+MT and DS+MT treatments received melatonin sprays every 3 days, while the other treatments were sprayed with an equivalent amount of distilled water. The concentration of 100 μmol/L melatonin was determined based on the preliminary research results of the research group(Yang et al., 2023).

### Measurement of above-ground agronomic traits

On days 0, 6, 12, and 18 after treatment, plant height, stem diameter, and leaf area were measured, with five biological replicates for each measurement. Plant height was defined as the distance from the cotyledon node to the main shoot apex. Stem diameter was measured using a caliper at a point 1 cm above the cotyledon node. Leaf area was calculated using the length-width coefficient method (0.75).

### Measurement of leaf relative water content

On days 0, 6, 12, and 18 after treatment, the relative water content of the leaves was measured using seedlings with inverted trifoliate leaves, with three biological replicates for each measurement. The relative water content was determined using the saturation-drying weighing method. In brief, the third functional leaf from the top of the main stem was collected and immediately weighed on an analytical balance to obtain the fresh weight (g). After recording this weight, the leaf was placed in deionized water and kept in a refrigerator at 4 °C for 8 to 12 hours to determine the turgid weight (g). Subsequently, the samples were dried in an oven at 70 °C for 72 hours, and the dry weight (g) of the leaf samples was measured. The calculation formula is: Relative Water Content = (Fresh Weight - Dry Weight) / (Turgid Weight - Dry Weight) (Yang et al., 2022).

### Analysis of root traits

On days 0, 6, 12, and 18 after treatment, total root length, total root surface area, and total root volume were measured, with three biological replicates for each measurement. The roots were placed without overlapping in a transparent acrylic box with a water depth of 1 cm for scanning (Epson V700; Epson, Suwa, Japan) to obtain root images. The total root length, surface area, and volume were analyzed using WinRHIZO software (WinRHIZO REG 2009, Canada).

### Measurement of biomass

On days 0, 6, 12, and 18 after treatment, the biomass of both the root system and the aerial parts was measured, with three biological replicates for each measurement. The seedlings were harvested, and the roots and aerial parts were separated and placed in bags. They were then heated at 105 °C for 30 minutes to inactivate enzymes and dried at 80 °C until reaching a constant weight to determine the biomass of the roots and aerial parts.

### Measurement of water and nutrient absorption rates

On day 18 after treatment, the specific root length absorption rates of water, nitrate, ammonium, and phosphate in the seedlings were measured, with three biological replicates for each measurement. The seedlings were preconditioned by transferring them to nutrient solution lacking macroelements for 48 hours prior to measurement. Twelve conical flasks containing 260 ml of normal nutrient solution were prepared, and 2 ml of the solution was collected as a sample before transferring the seedlings. After the seedlings were transferred to the flasks for 4 hours, the remaining volume of nutrient solution was measured, and the water absorption rate was calculated based on the difference in nutrient concentration before and after absorption. Additionally, 2 ml samples of the remaining nutrient solution were collected and stored at −20 °C. Root images were captured at the end of sampling for total root length analysis. A continuous flow chemical analyzer (SmartChem 600, France) was used to determine the concentrations of nitrate, ammonium, and phosphate in the water samples, and the nutrient absorption rates were calculated based on the difference in nutrient concentration before and after absorption (Roy et al., 2021).

### Analysis of root cortex anatomy

On days 0, 6, 12, and 18 after treatment, the root cortex cells were examined. Lateral roots were selected at a position 1 cm from the base of the main root, with a lateral root length of approximately 20 cm. Root segments were taken from the lateral roots at 1 cm and 13 cm from the root tip (0.5 cm segments), with three biological replicates and three technical replicates for each measurement (Fig. 1). The collected root segments were immediately placed in FAA fixative and fixed at 4 °C for 24 hours. Using the paraffin sectioning technique, the roots were subjected to dehydration, clarification, embedding, and sectioning to produce anatomical slides. After staining with acridine orange (Henry & Deacon, 1981), images of the sections were captured using a fluorescence microscope (Axio Imager M2, Zeiss, Germany). The root cross-sectional area, stele area, and cortical thickness were measured using ImageJ software, and the area of the living cortex was calculated as (root cross-sectional area - stele area) (Jaramillo et al., 2013).

**Figure 1.**
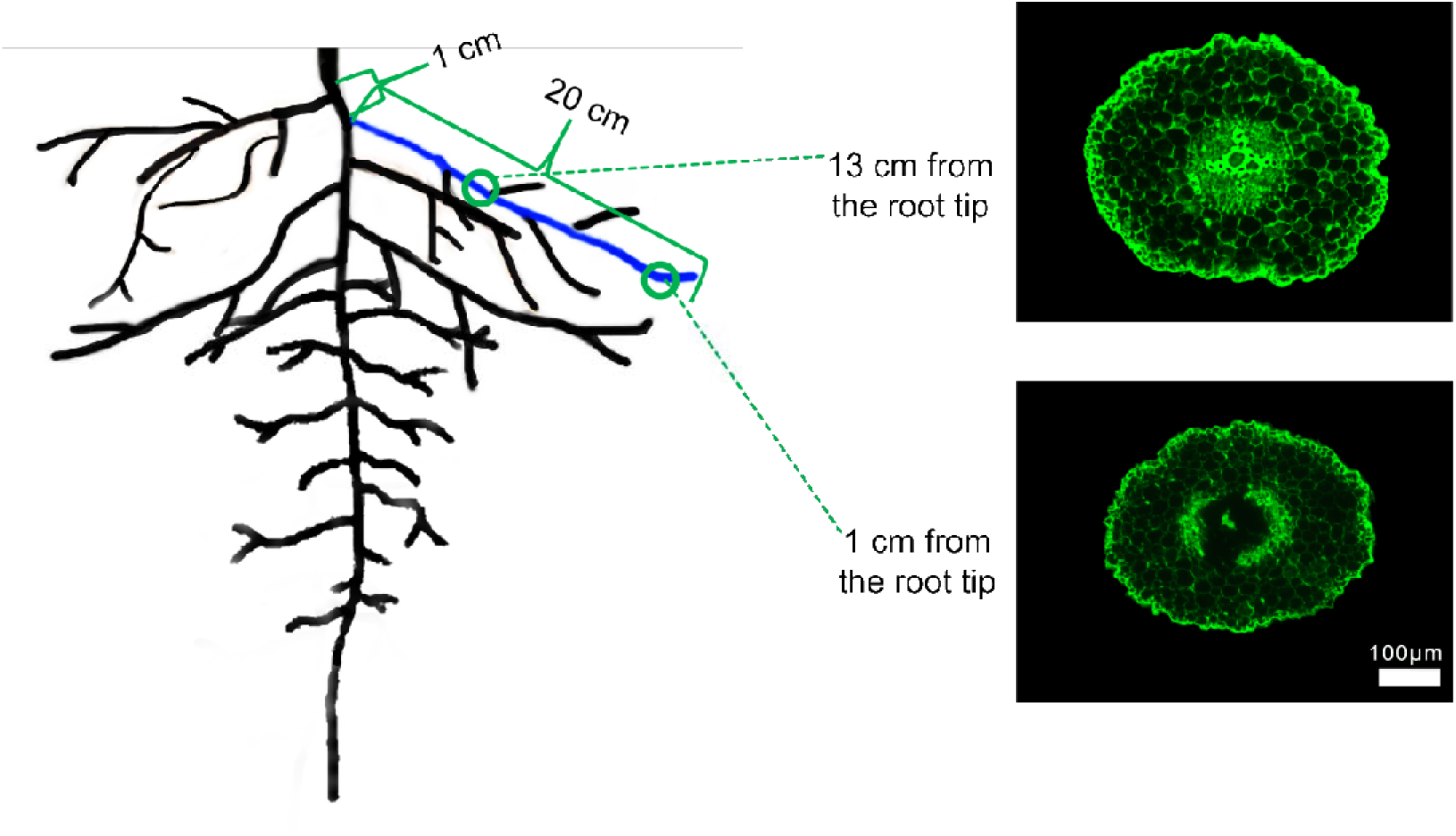
Schematic diagram of sampling for root cortex anatomy. Note: 1 cm represents the distance from the lateral root growth point to the base of the main root. 20 cm is the length of the sampled lateral root. The blue part indicates the selected lateral root. The green rings mark the positions for taking 0.5 cm root segments at different distances from the root tip.

### Measurement of osmotic regulatory substances

On days 0, 6, 12, and 18 after treatment, the content of soluble sugar and soluble protein were measured, with three biological replicates for each measurement. In brief, 0.2 g of plant roots was weighed, rapidly frozen in liquid nitrogen, and stored at −80 °C. The content of soluble sugars was determined using the anthrone colorimetric method, while the content of soluble protein was assessed using the Coomassie Brilliant Blue method (Jiancheng Biotechnology, Nanjing, China).

### Measurement of Root Respiration Rate and Activity of Respiratory Metabolism Enzymes

On days 0, 6, 12, and 18 after treatment, the root respiration rate and the activity of respiratory metabolism enzymes in the roots were measured, with three biological replicates for each measurement. In brief, a Li-840A CO_2_ analyzer was used for measurement. Clean roots were quickly dried of surface moisture using absorbent paper and immediately placed into a custom-made 18 ml respiration chamber. The gas circuit was maintained as closed throughout the measurement process, and measurements were completed when the curve stabilized. The roots were then collected for total root length measurement, and the specific root length respiration rate was calculated. The activities of phosphofructokinase, glucose-6-phosphate dehydrogenase, and malate dehydrogenase in roots were determined using the kits provided by Abbkine (Scientific Co., Ltd, Wuhan, China). Sampling was conducted according to “Measurement of osmotic regulatory substances”, and the determination was carried out in accordance with the kit instructions.

### Statistical analysis

Data recording and organization were performed using Microsoft Excel 2010 (Microsoft Corporation, Redmond, WA, USA). One-way ANOVA was conducted using IBM SPSS Statistics 27.0 (IBM Corp, Armonk, NY, USA), and the Duancan method was used for multiple comparisons and significance tests, with a significance level of *p* < 0.05. Graphs were created using GraphPad Prism 9.0 (GraphPad Software, Inc., San Diego, CA, USA), and correlation analysis charts were plotted using Origin 2021 (OriginLab Corporation, Northampton, MA, USA).

## Results

### Effects of exogenous melatonin on the shoot growth of cotton seedlings under drought stress

Melatonin significantly improved plant height, stem diameter, leaf area, shoot biomass, and leaf water content under drought stress (Fig. 2). Compared with the control, after 18 days of drought stress treatment, the plant height, stem diameter, leaf area, shoot biomass, and relative water content of leaves decreased by 44.73%, 47.20%, 82.39%, 74.06%, and 44.40%, respectively (*p* < 0.05). However, exogenous application of melatonin could promote the growth and development of cotton, especially under drought stress conditions. Specifically, compared with the well water treatment, the stem diameter and shoot biomass of the well water and MT treatment increased significantly by 10.53% and 15.72%, respectively. Meanwhile, the drought stress and MT treatment increased these parameters by 9.35%, 24.15%, 58.49%, 40.63%, and 57.56%, respectively (*p* < 0.05). In conclusion, exogenous application of melatonin can significantly reduce the adverse effects of drought stress on shoot traits, with an obvious alleviating effect.

**Figure 2.**
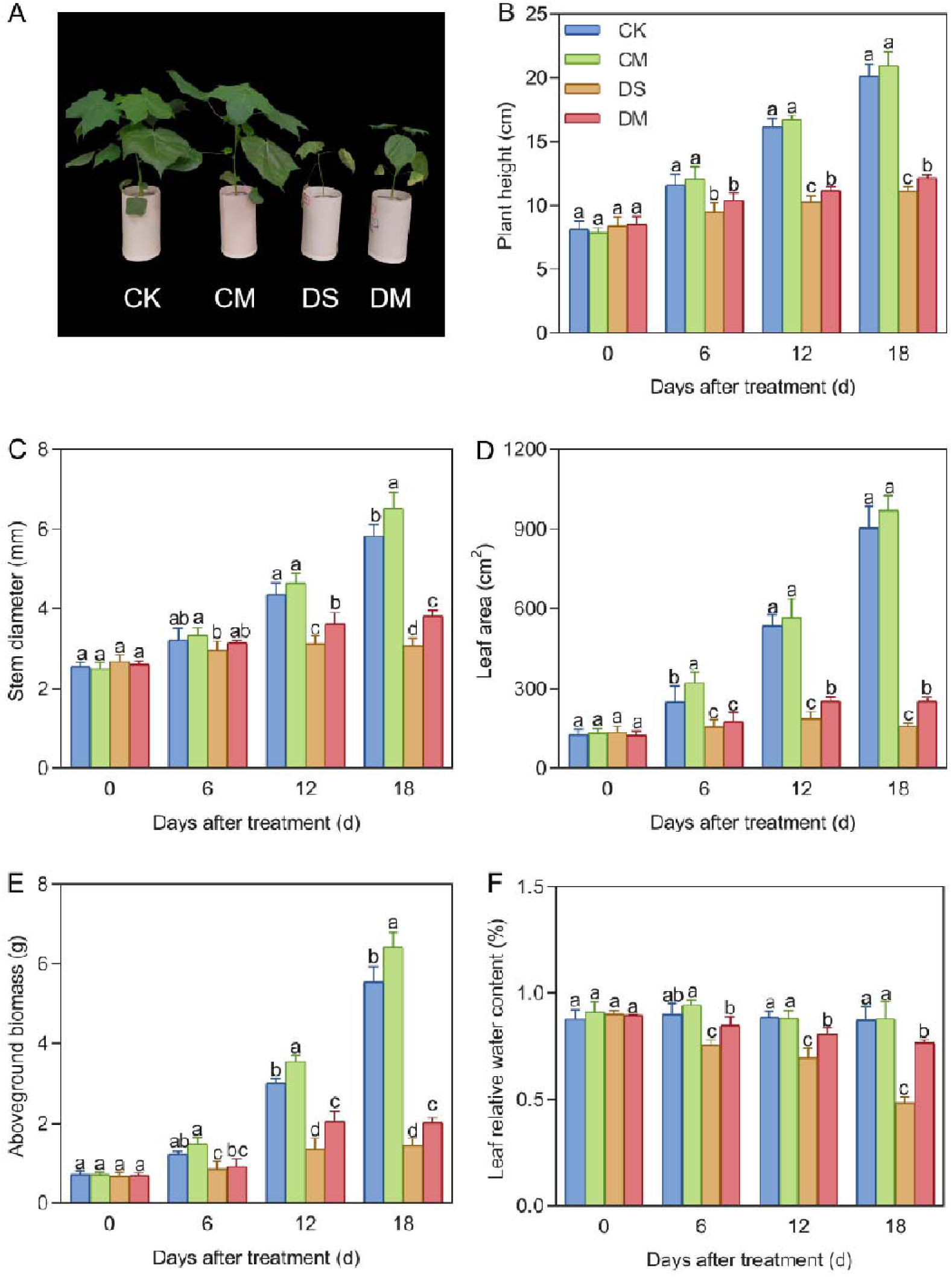
Effects of different treatments on above-ground indicators. (A) Phenotype of cotton 18 days after treatment, (B) Plant height, (C) Stem diameter, (D) Leaf area, (E) Above-ground biomass, and (F) Leaf relative water content. CK, well water; CM, well water and melatonin; DS, drought stress; DM, drought stress and melatonin. Different lowercase letters indicate significant differences among different treatments at the same time period (*p* < 0.05). (B-D): Values are means ± SE (n=5), (E-F): Values are means ± SE (n=3).

### Effects of exogenous melatonin on the growth and development of cotton roots under drought stress

Melatonin enhanced total root length, surface area, volume, and dry weight under drought stress (Fig. 3). Compared with the control, after 18 days of drought stress treatment, the total root length, total root surface area, total root volume, and root dry weight decreased by 71.96%, 73.36%, 74.58%, and 65.45%, respectively (*p* < 0.05). However, exogenous application of melatonin could promote the growth and development of roots, especially under drought-stress conditions. Specifically, compared with the well water treatment, the total root surface area, total root volume, and root dry weight of the well water and MT treatment increased significantly by 10.71%, 15.01%, and 20.72%, respectively. Meanwhile, the total root length, total root surface area, total root volume, and root dry weight of the drought stress and MT treatment increased significantly by 52.90%, 58.36%, 63.56%, and 40.26%, respectively. In conclusion, exogenous application of melatonin can alleviate the growth inhibition of cotton roots caused by drought stress and play a positive role in promoting the growth of cotton roots.

**Figure 3.**
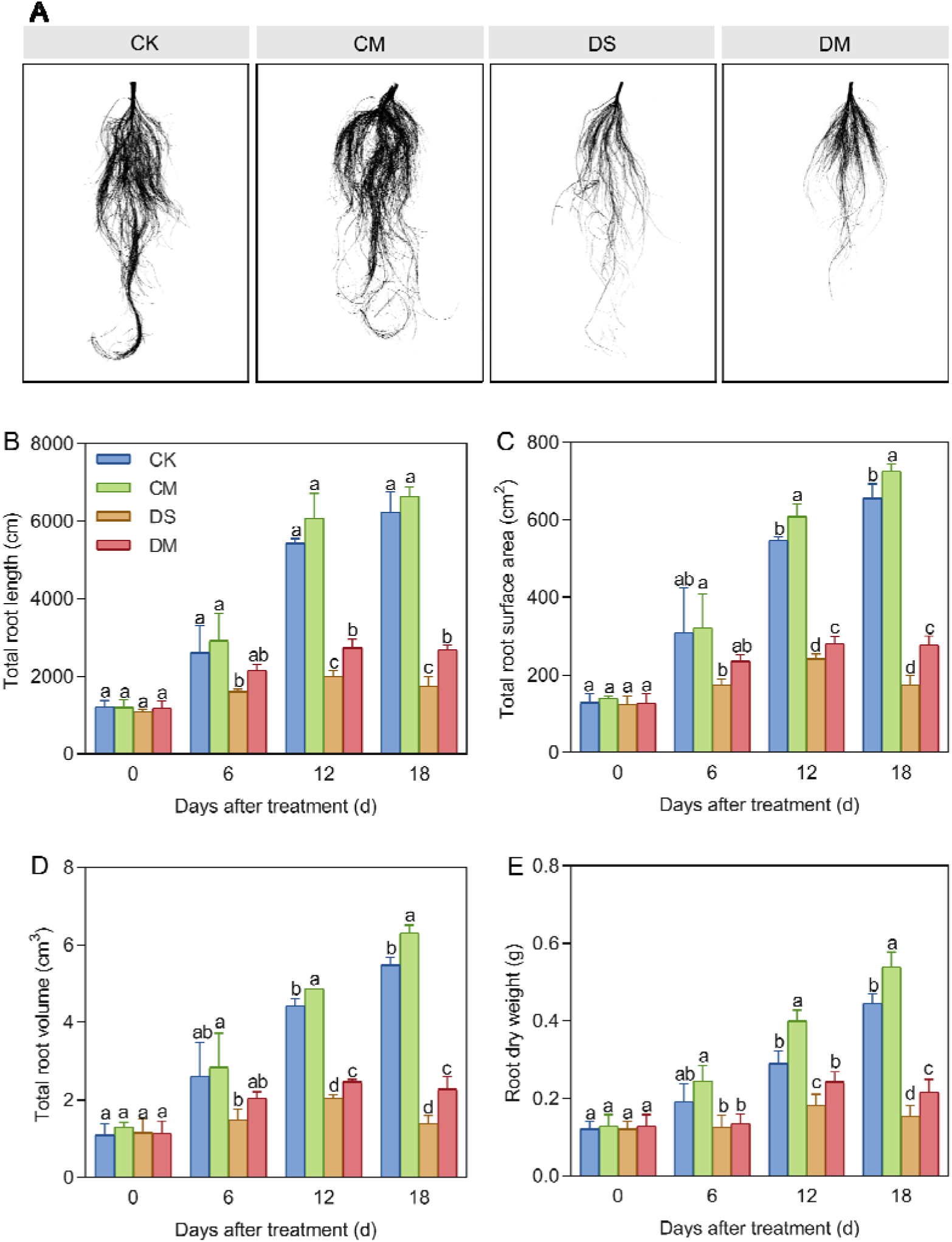
Effects of different treatments on root morphological indices. (A) Root phenotype of cotton 18 days after treatment, (B) Total root length, (C) Total root surface area, (D) Total root volume, and (E) Root dry weight. CK, well water; CM, well water and melatonin; DS, drought stress; DM, drought stress and melatonin. Different lowercase letters indicate significant differences among different treatments at the same time period (*p* < 0.05). Values are means ± SE (n=3).

### Effects of exogenous melatonin on the absorption rates of water and nutrients in cotton roots under drought stress

Melatonin significantly promoted the absorption rates of water, nitrate, ammonium, and phosphate in cotton roots under drought-stress treatment (Fig. 4). Compared with the control, after 18 days of drought stress treatment, the specific root length absorption rates of water, nitrate, ammonium, and phosphate decreased by 60.25%, 54.40%, 59.60%, and 67.01%, respectively (*p* < 0.05). However, exogenous application of melatonin could enhance the absorption capacity of roots, especially under drought-stress conditions. Specifically, compared with the well water treatment, the specific root length absorption rates of water and nitrate in the well water and MT treatment increased significantly by 11.85% and 16.89%, respectively. Meanwhile, in the drought stress and MT treatment, the specific root length absorption rates of water, nitrate, ammonium, and phosphate increased significantly by 61.97%, 49.52%, 59.21%, and 83.26%, respectively (*p* < 0.05).

**Figure 4.**
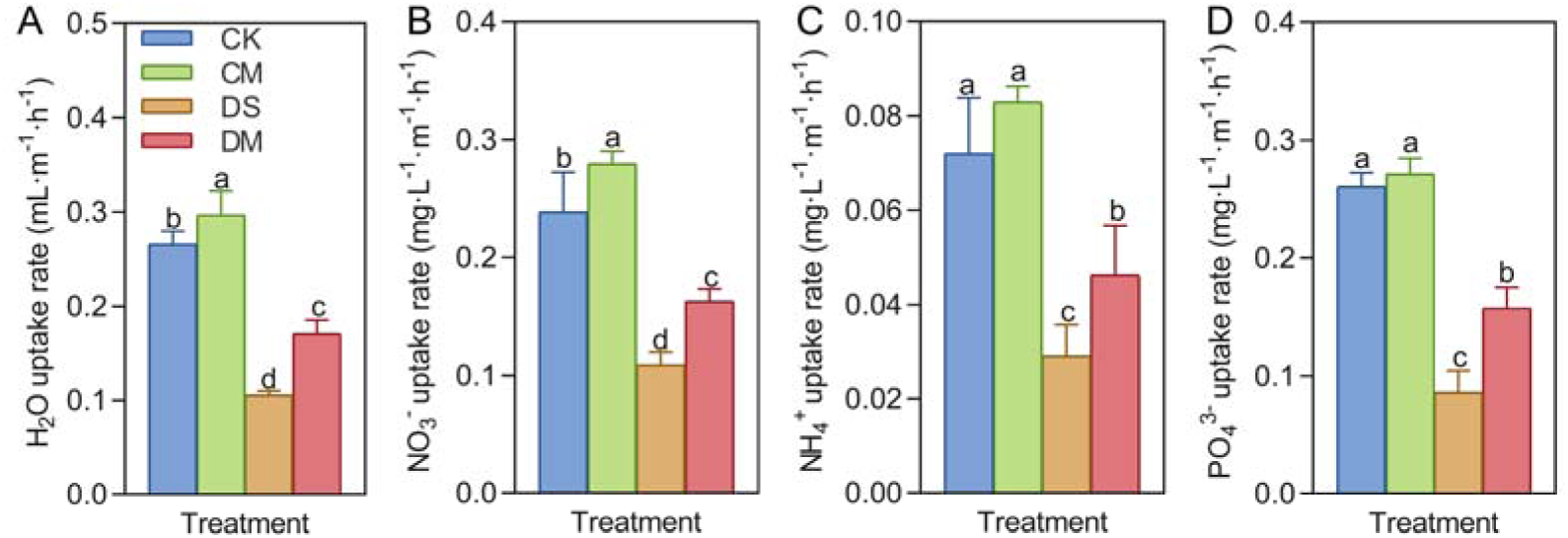
Effects of different treatments on the specific root length absorption rate 18 days after treatment. (A) Water uptake rate, (B) Nitrate uptake rate, (C) Ammonium salt uptake rate, and (D) Phosphate uptake rate. CK, well water; CM, well water and melatonin; DS, drought stress; DM, drought stress and melatonin. Different lowercase letters indicate significant differences among different treatments in the same period (*p* < 0.05). Values are means ± SE (n=3).

### Effects of exogenous melatonin on the root cortex of cotton seedlings under drought stress

Melatonin significantly promoted the living cortical area and cortical thickness of cotton roots under drought-stress treatment (Figs. 5 and 6). The data showed that at a position 1 cm from the root tip, Compared with the control, after 18 days of drought stress treatment, the living cortical area and cortical thickness of the roots decreased by 54.32% and 36.00%, respectively (*p* < 0.05) (Figs. 6A,C). However, exogenous application of melatonin could promote the living cortical area of cotton roots under drought stress. Specifically, the living cortical area in the drought stress and MT treatment was significantly increased by 26.34% compared with the drought stress treatment (Fig. 6A). Meanwhile, at a position 13 cm from the root tip, significant differences began to appear on the 12th day. Compared with the well water, after 18 days of drought stress treatment, the living cortical area and cortical thickness of the roots decreased by 45.50% and 28.19%, respectively (*p* < 0.05), while the living cortical area of the roots in the well water and MT treatment was significantly increased by 9.69%. At the same time, compared with the drought stress treatment, the living cortical area and cortical thickness of the roots in the drought stress and MT treatment were significantly increased by 59.00% and 21.24%, respectively (Figs. 6B,D). In conclusion, exogenous application of melatonin can maintain the activity of root cortex cells in cotton under drought stress and improve root adaptability.

**Figure 5.**
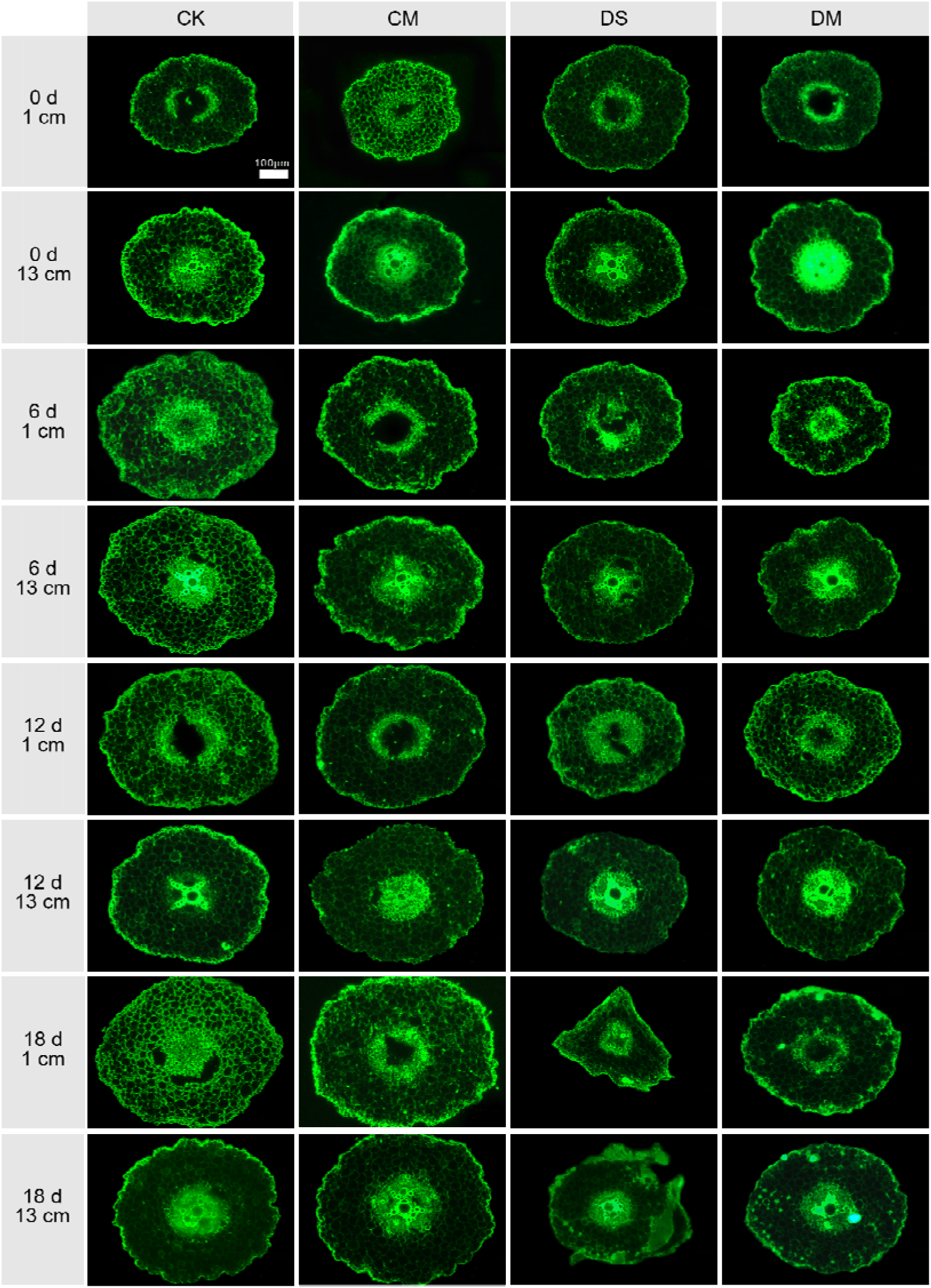
Effects of different treatments on the root cross-sections at 1 cm and 13 cm from the root tip at 0 d, 6 d, 12 d and 18 d after treatment. CK, well water; CM, well water and melatonin; DS, drought stress; DM, drought stress and melatonin. Scale bar: 100 μm

**Figure 6.**
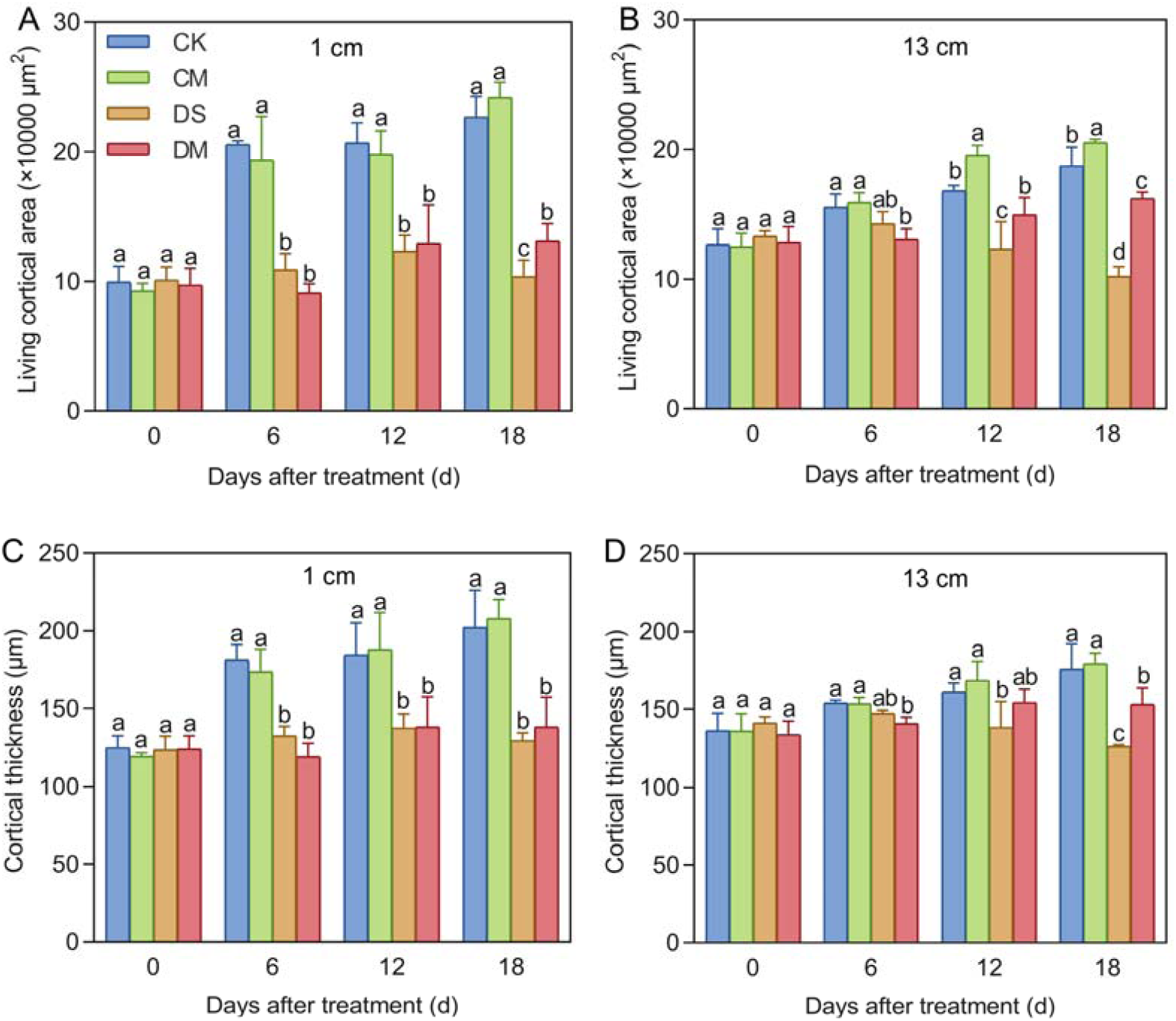
Effects of different treatments on root cortex cells. (A) Living cortical area at 1 cm from the root tip, (B) Living cortical area at 13 cm from the root tip, (C) Cortical thickness at 1 cm from the root tip, and (D) Cortical thickness at 13 cm from the root tip. CK, well water; CM, well water and melatonin; DS, drought stress; DM, drought stress and melatonin. Different lowercase letters indicate significant differences among different treatments at the same time period (*p* < 0.05). Values are means ± SE (n=3).

### Effects of exogenous melatonin on the contents of osmotic regulatory substances in the roots of cotton seedlings under drought stress

Melatonin significantly promoted the accumulation of soluble sugar and soluble protein in cotton seedlings under drought-stress treatment (Fig. 7). Compared with the control, after 18 days of drought stress treatment, the content of soluble sugars increased by 14.31% (*p* < 0.05), while the content of soluble protein decreased significantly by 12.88%. However, exogenous application of melatonin could effectively increase the osmotic regulatory substances. Specifically, compared with the well water treatment, the content of soluble sugars in the well water and MT treatment increased significantly by 12.37%. Meanwhile, in the drought stress and MT treatment, the content of soluble sugar and soluble protein increased significantly by 11.90% and 16.75%, respectively (*p* < 0.05). In conclusion, exogenous application of melatonin plays a positive role in promoting the increase in the content of soluble sugar and soluble protein under drought stress, and can effectively alleviate the damage to roots caused by drought stress.

**Figure 7.**
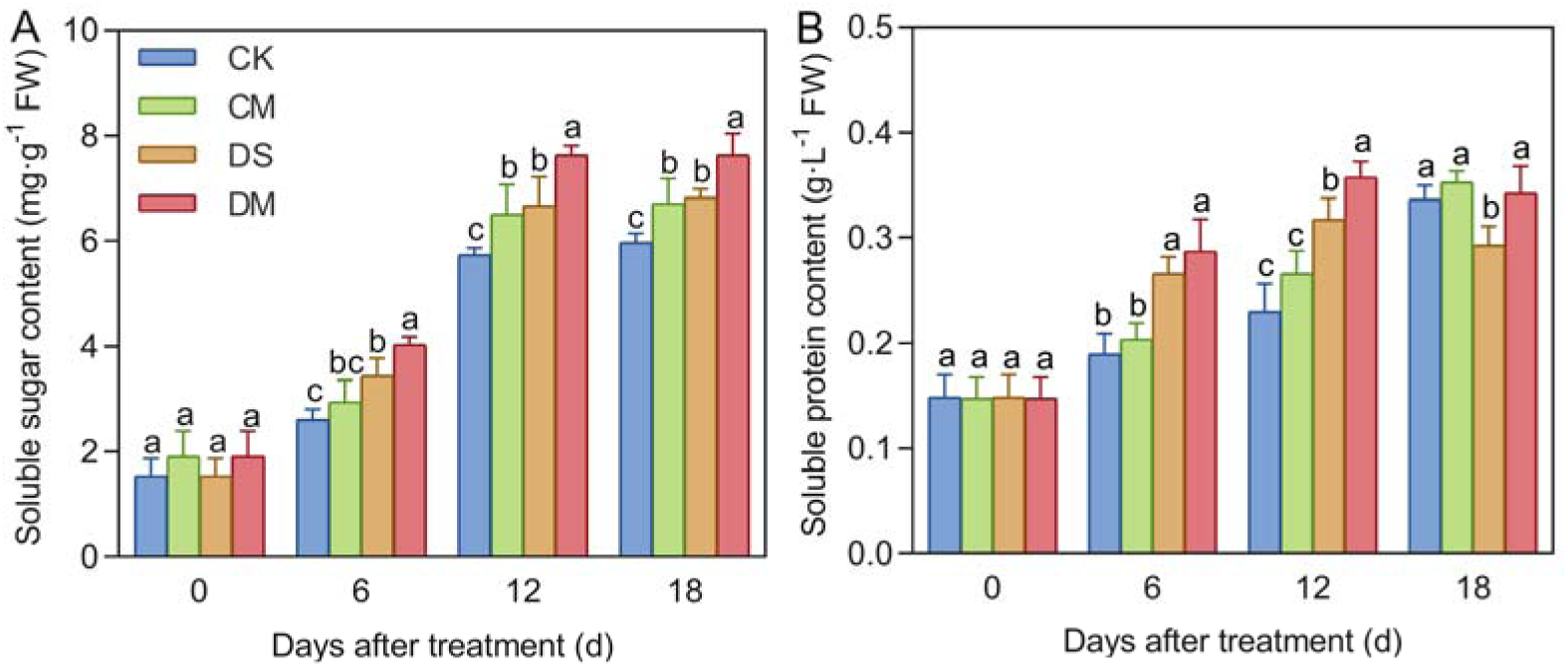
Effects of different treatments on osmotic regulatory substances in roots. (A) Soluble sugar content, and (B) Soluble protein content. CK, well water; CM, well water and melatonin; DS, drought stress; DM, drought stress and melatonin. Different lowercase letters indicate significant differences among different treatments at the same time period (*p* < 0.05). Values are means ± SE (n=3).

### Effects of exogenous melatonin on root metabolism of cotton seedlings under drought stress

Melatonin significantly enhanced the activities of root respiratory metabolic enzymes and the root respiration rate in cotton seedlings under drought-stress treatment (Fig. 8). Compared with the control, after 18 days of drought stress treatment, the activities of phosphofructokinase, glucose-6-phosphate dehydrogenase, malate dehydrogenase, and the specific root length respiration rate decreased by 25.16%, 47.50%, 75.37%, and 73.87%, respectively (*p* < 0.05). However, exogenous application of melatonin could significantly improve the normal level of root metabolism and mitigate the adverse effects of drought treatment. Specifically, compared with the well water treatment, the activities of glucose-6-phosphate dehydrogenase, malate dehydrogenase, and the specific root length respiration rate in the well water and MT treatment increased significantly by 20.73%, 10.45%, and 12.38%, respectively. Meanwhile, in the drought stress and MT treatment, the activities of phosphofructokinase, glucose-6-phosphate dehydrogenase, malate dehydrogenase, and the specific root length respiration rate increased significantly by 14.29%, 30.23%, 48.47%, and 41.06%, respectively (*p* < 0.05).

**Figure 8.**
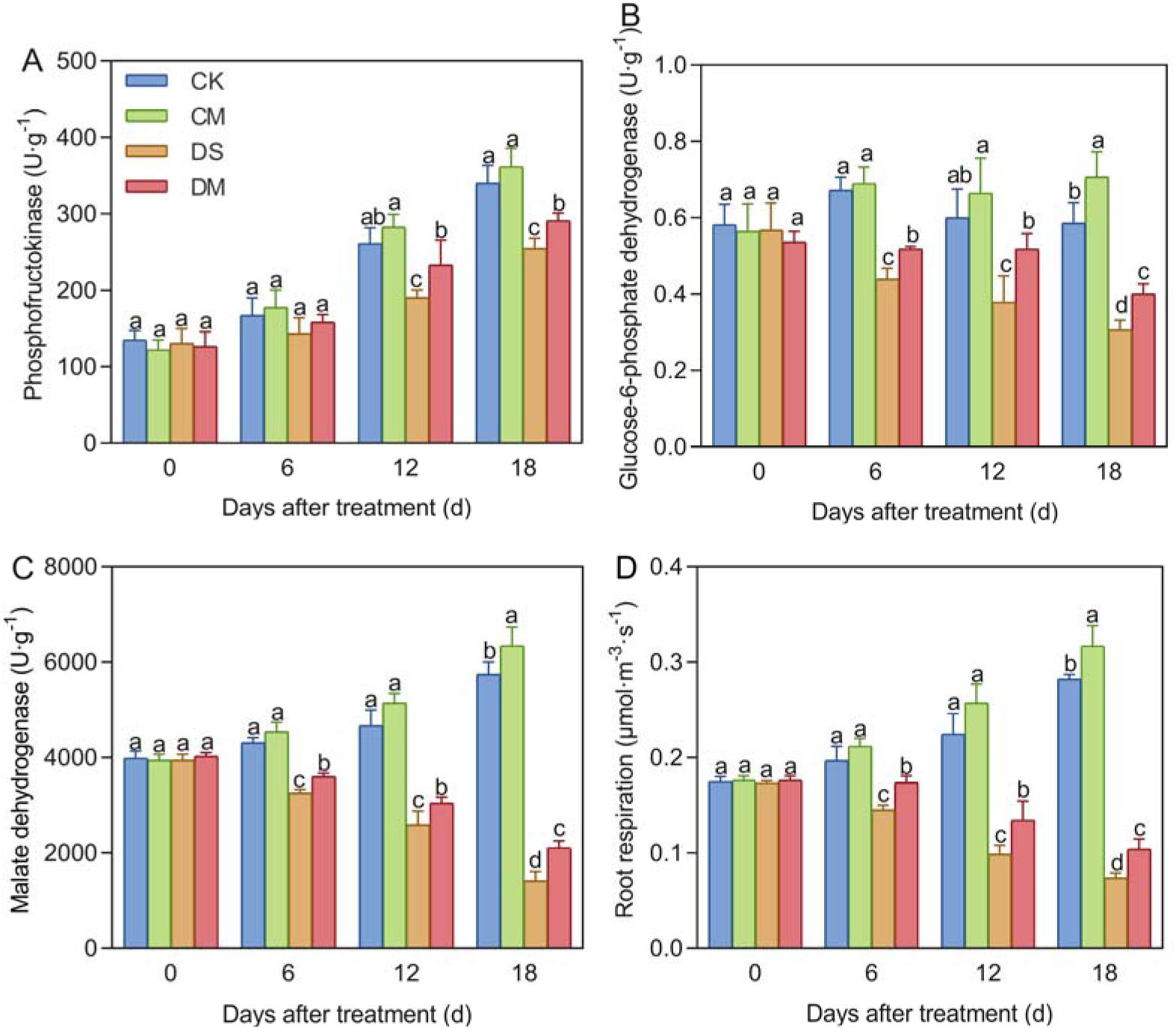
Effects of different treatments on root metabolism. (A) Phosphofructokinase, (B) Glucose-6-phosphate dehydrogenase, (C) Malate dehydrogenase, and (D) Specific root length respiration rate. CK, well water; CM, well water and melatonin; DS, drought stress; DM, drought stress and melatonin. Different lowercase letters indicate significant differences among different treatments at the same time point (*p* < 0.05). Values are means ± SE (n=3).

### Correlation analysis

As shown in Fig. 9A, under normal environmental conditions, the active root cortical area was significantly positively correlated with soluble protein and positively correlated with soluble sugars. This indicates that there is a significant positive correlation between root cortical cells and root osmotic adjustment under normal environmental conditions.

**Figure 9.**
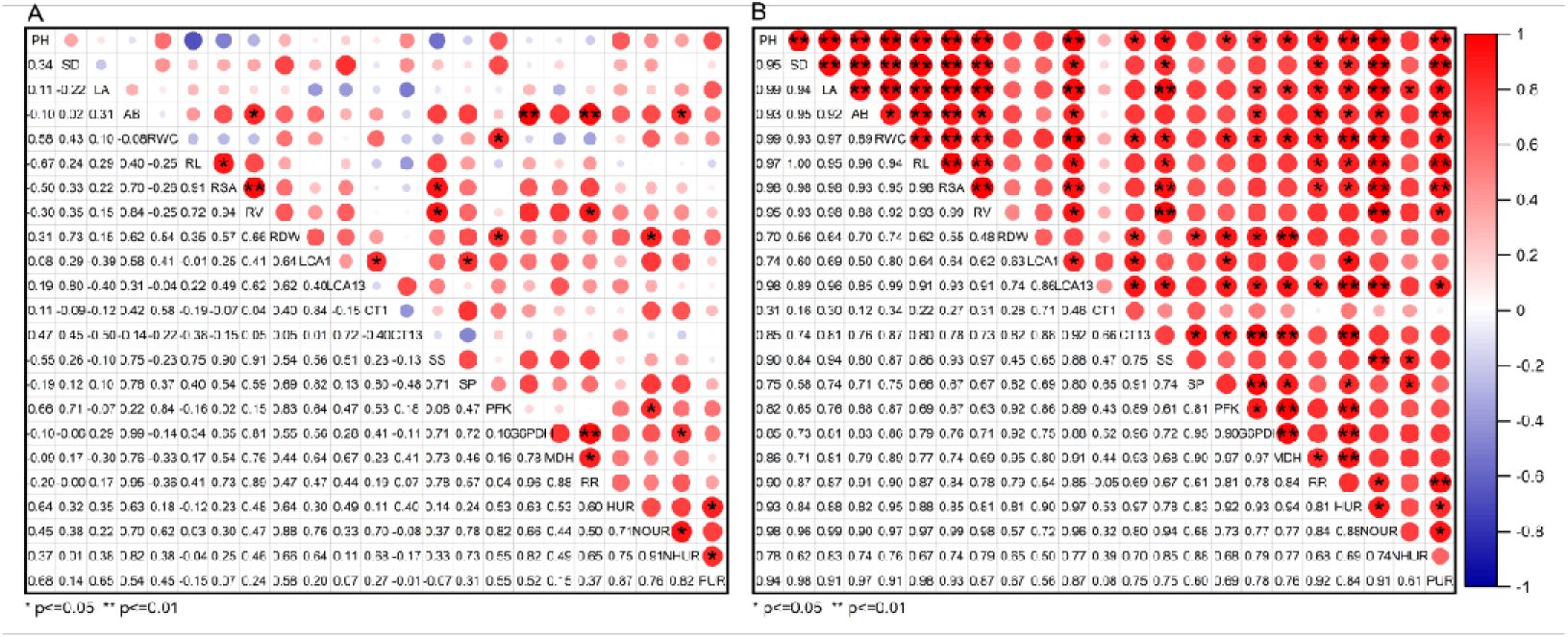
Correlation analysis of traits of cotton seedlings under well water (A) and drought stress (B) treatments with the application of melatonin. Note: PH: Plant height, SD: Stem diameter, LA: Leaf area, AB: Aboveground biomass, RWC: Leaf relative water content, RL: Total root length, RSA: Total root surface area, RV: Total root volume, RDW: Root dry weight, LCA1: Living cortical area at 1 cm from the root tip, LCA13: Living cortical area at 13 cm from the root tip, CT1: Cortical thickness at 1 cm from the root tip, CT13: Cortical thickness at 13 cm from the root tip, SS: Soluble sugar content, SP: Soluble protein content, PFK: Phosphofructokinase, G6PDH: Glucose-6-phosphate dehydrogenase, MDH: Malate dehydrogenase, RR: Specific root length respiration rate, HUR: Specific root length Water uptake rate, NOUR: Specific root length Nitrate uptake rate, NHUR: Specific root length Ammonium uptake rate, PUR: Specific root length Phosphate uptake rate.

As shown in Fig. 9B, under drought stress, the active root cortical area was significantly positively correlated with soluble sugars, and the root cortical thickness was significantly positively correlated with soluble protein. This indicates that when melatonin is applied under drought stress, there is a significant positive correlation between root cortical cells and root osmotic adjustment. The active root cortical area was significantly positively correlated with phosphofructokinase, glucose-6-phosphate dehydrogenase, malate dehydrogenase, and the specific root length respiration rate. Therefore, when melatonin is applied under drought stress, there is a significant positive correlation between root cortical cells and root respiratory metabolism. The living cortical area of roots was significantly positively correlated with the specific root length absorption rates of water, nitrate, and phosphate, and strongly positively correlated with the specific root length absorption rate of ammonium. The root cortical thickness was also positively correlated with the specific root length absorption rate of ammonium. Therefore, when melatonin is applied under drought stress, there is a significant positive correlation between root cortical cells and root absorption function. The active root cortical area was significantly positively correlated with the total root length, total root surface area, and total root volume of the roots, and significantly positively correlated with the plant height, stem diameter, leaf area, shoot biomass, and relative leaf water content of the above-ground parts. The root cortical thickness was significantly positively correlated with the root dry weight. Therefore, when melatonin is applied under drought stress, there is a significant positive correlation between root cortical cells and the growth status of cotton. In conclusion, exogenous melatonin promotes root growth by increasing the area of root cortical cells under drought stress, facilitating osmotic accumulation, enhancing respiratory metabolism, and improving the root absorption rate, ultimately alleviating the damage caused by drought stress.

## Discussion

### Exogenous melatonin maintains the activity of root cortical cells under drought stress

In the research field of plants’ response to drought stress, predecessors have conducted in-depth studies on the root cortical structure. Numerous studies have shown that the structure undergoes significant changes under drought stress. Specifically, drought stress causes mechanical damage to the cortical cells of fine roots in grapevines, such as the formation of lacunae, resulting in permanent structural damage to root cortical cells and a reduction in the active root cortical area (Cuneo et al., 2016). Moreover, a phenomenon of cortical cell shedding, known as root cortex death, has been observed in wheat and barley. This shedding is accelerated in water-deficient environments, leading to a decrease in cortical thickness, the disintegration of cortical cells, and the redistribution of remaining nutrients to rapidly growing root tissues (Liljeroth, 1995; Schneider et al., 2017). These results are similar to those of this study on cotton crops. That is, under drought stress, the active root cortical area and cortical thickness gradually decrease with the increase in drought stress duration (Figs. 5 and 6). This indicates that the impact of drought stress on the root cortical structure of plants is somewhat universal and can damage root cortical cells.

Melatonin is a natural indole molecule widely present in animals and plants. Exogenous application of melatonin has been proven to be an effective way to enhance the drought resistance of plants in crop production (Dubbels et al., 1995; Guo et al., 2023). This study found that under drought stress, exogenous application of melatonin can alleviate the growth inhibition of cotton roots caused by drought stress and promote total root length, total root surface area, total root volume, and root dry weight (Fig. 3). However, under severe abiotic stress, for crops to maintain growth, roots need to have sufficient energy, and the active root cortical area must be maintained at a certain level; neither can be lacking (Du et al., 2024). Therefore, this experiment explored the effect of exogenous melatonin application on the root cortex under drought stress. It was also found that exogenous application of melatonin can increase the area of cotton root cortical cells under drought stress and improve the drought tolerance of cotton (Fig. 6). This is similar to previous research results. Under salt stress, melatonin accumulates significantly in the root cortical cells of sunflowers, and exogenous melatonin can regulate the root growth of sunflower seedlings, indicating that there is an association between the accumulation of melatonin in root cortical cells and the regulation of root growth (Mukherjee et al., 2014).

There are some key points in the sampling process of the anatomical structure of cotton roots that deserve in-depth discussion. Cotton is a tap-rooted plant with a main root and lateral roots. Lateral roots were ultimately selected for the study, mainly because the physiological activities of lateral roots may be more active, they account for a larger proportion of the root system, and their physiological state and structural characteristics can better reflect the dynamic changes of cotton roots during the overall growth process. At the same time, there is no difference in the response of the anatomical structures of the two types of roots to drought treatment among root types (Jaramillo et al., 2013). Since the roots are constantly growing and developing, to effectively reduce experimental errors, the sampling location of the roots was strictly standardized during sampling (Sidhu et al., 2024). Specifically, lateral roots about 20 cm in length, 1 cm away from the base of the main root, were selected. Then, root segments (0.5 cm in length) at 1 cm and 13 cm from the root tip were precisely measured and cut (Fig. 1). Through this highly unified sampling method, every effort was made to ensure that the obtained samples had a high degree of consistency in physiological state and structural characteristics, thereby improving the accuracy and reliability of the experimental data.

### Exogenous melatonin promotes the accumulation of osmotic regulatory substances, enhances root metabolism, and improves root absorption function under drought stress

Under adverse conditions, the osmotic balance within plant cells is disrupted. To maintain cell turgor pressure and enhance their adaptability to unfavorable environments, plants increase the content of osmotic regulatory substances (Yancey, 2005). Exogenous melatonin can promote the synthesis of osmotic regulatory substances by regulating the expression of related genes. Previous studies have shown that after melatonin treatment, the expression levels of genes related to soluble sugar synthesis in early-maturing peach fruit cells are upregulated, leading to an increase in the soluble sugar content (Zhou et al., 2023). This is similar to the results of this study on cotton crops. Specifically, the content of soluble sugar and soluble protein in cotton roots were significantly higher than those of the control 6 days after drought-stress treatment, and exogenous application of melatonin significantly promoted the accumulation of soluble sugar and soluble protein in the roots of cotton seedlings under drought stress (Fig. 7). The possible reason why the content of soluble protein under drought stress was lower than that of the control treatment on the 18th day after treatment was the severity of drought stress (Fig. 7). The above results indicate that exogenous melatonin plays a crucial role in stress resistance under drought stress by increasing the content of osmotic regulatory substances.

Drought stress significantly affects the respiratory metabolism pathways of plant roots. Specifically, in cotton roots, the activities of phosphofructokinase, glucose-6-phosphate dehydrogenase, and malate dehydrogenase decrease, which impacts the energy supply of plants (Liu et al., 2024). Meanwhile, through subjecting various plants to different degrees of drought treatment, it has been found that as the severity of drought increases and the duration prolongs, the root respiration rate of plants shows a downward trend (Fathi & Barari, 2016). A similar pattern emerged in this study under drought stress, that is, the activities of respiratory metabolic enzymes and the specific root length respiration rate of cotton roots decreased significantly 12 days after drought-stress treatment (Fig. 8). This indicates that drought stress can significantly affect root metabolism. In addition, a sharp decrease in root metabolic activities may limit crop growth and yield, and even lead to crop death (Du et al., 2024). However, exogenous melatonin can improve the root vitality of cotton under drought stress, yet whether its underlying mechanism is directly related to the increase in the root respiratory metabolism pathway remains to be further explored (Zhu et al., 2024). In this study, it was discovered for the first time that exogenous application of melatonin can significantly increase the activities of phosphofructokinase, glucose-6-phosphate dehydrogenase, and malate dehydrogenase in cotton roots under drought-stress treatment, and at the same time increase the root respiration rate (Fig. 8). This suggests that exogenous melatonin may enhance the stress resistance of cotton by activating these key enzymes, regulating the respiratory metabolism pathway, and providing more sufficient energy for the roots, which supports the role of melatonin in regulating root metabolism. This is consistent with the research results of Turk and Genisel (2020) on the growth of maize under cold stress, where melatonin enhanced the mitochondrial respiration rate and energy production. Phosphofructokinase is a key rate-limiting enzyme in the glycolysis process, and its activity may increase with the increase in drought-stress duration (Fig. 8), thus accelerating the decomposition of sugars such as glucose and providing more intermediate metabolites and energy for the roots (Dwivedi et al., 2023).

Under stress conditions, the absorption and utilization of water and nutrients in plants are often severely disrupted. Melatonin exhibits a positive regulatory function. It can reduce the oxidative damage to roots and the entire plant caused by stress by enhancing the plant’s antioxidant defense system, maintaining the normal physiological functions of roots, and thus minimizing the impact on root absorption (Sun et al., 2022). The positive effect of melatonin in alleviating drought in maize has been widely reported. PEG-induced water shortage significantly decreases the root hydraulic conductivity, affecting the root’s ability to absorb water and transport it to the above-ground parts. However, after the application of melatonin, the root hydraulic conductivity is significantly enhanced, improving the root’s water absorption and transportation functions (Qiao et al., 2020). Similarly, there are reports on apples. Under drought stress, the absorption fluxes of nitrogen, phosphorus, and potassium by apple roots decrease significantly. But after long-term exogenous application of melatonin, the absorption of nitrogen by roots is promoted by regulating the gene expression and activity of root-related nitrogen transporters, and the absorption of phosphorus by roots is enhanced by accelerating the secretion of acid phosphatase by roots (Liang et al., 2018). This is consistent with the results of this study. This study found that exogenous melatonin significantly promotes the absorption rates of water, nitrate, ammonium, and phosphate by cotton roots under drought stress (Fig. 4). Therefore, melatonin improves the water and nutrient use efficiency of plants under drought stress by regulating the absorption of root-related nutrients, thereby ensuring the normal physiological functions of roots under stress conditions. In summary, exogenous melatonin increases the active root cortical area, promotes osmotic accumulation, enhances respiratory metabolism, improves the absorption of water and nutrients, and thus promotes root growth.

### Exogenous Melatonin Improves the Drought Resistance of Cotton under Drought Stress

This study found that exogenous melatonin increases the active root cortical area of cotton, promotes the accumulation of osmotic regulatory substances, enhances root respiratory metabolism, and improves the root absorption rate, ultimately improving the drought resistance of cotton under drought stress. The specific mechanisms by which exogenous melatonin improves the above-mentioned root-related indicators and physiological processes to enhance the drought resistance of cotton mainly include three aspects:

First, an increase in the active root cortical area enlarges the contact area between the roots and the soil. Root hairs are structures formed by the outward protrusion of epidermal cells. Increasing the active root cortical area also increases the number of root hairs, leading to increased water absorption (Tanaka et al., 2014). Moreover, an intact active root cortex can provide more extensive intercellular connections and material transport channels, which is conducive to the absorption, transportation, and storage of water and nutrients in the soil by the roots, enhancing the plant’s survival ability in adverse environments (Cuneo et al., 2016). The increase in the active root cortical area provides more space for the accumulation of osmotic regulatory substances (Xiao et al., 2023). At the same time, it also increases basic root metabolic consumption, but a larger active root cortical area also provides more space for the distribution of mitochondria (Lambers et al., 2006). Therefore, increasing the active root cortical area means improving the drought resistance of cotton.

Second, the accumulation of osmotic regulatory substances in cells can rapidly decrease the cell water potential, causing water to enter the cells from the soil along the water potential gradient, maintaining the turgor pressure of the cells, and ensuring the normal physiological functions of the cells (Ozturk et al., 2021). Osmotic regulatory substances can also protect the cell’s biomembrane, prevent membrane lipid peroxidation, and ensure the normal metabolism and material transport of the cells, thus directly improving the drought resistance of the plant (Singh et al., 1972; Tian et al., 2013). The accumulation of substances requires energy consumption. For example, the synthesis of proline requires the participation of ATP and reducing power (NADPH) (Zhu et al., 2021). Third, enhanced root respiratory metabolism can provide energy for various physiological activities of the roots, maintain water balance, and a high rate of ion absorption and transportation, thereby promoting root growth (Griffiths et al., 2021). Therefore, promoting the accumulation of osmotic regulatory substances and enhancing root respiratory metabolism can improve the resistance of cotton to drought stress.

Finally, increasing the absorption rate of water and nutrients can directly provide the plant with sufficient water to meet its physiological needs in a drought environment. Rapid water absorption can ensure the normal progress of photosynthesis and transpiration in leaves, providing energy and raw materials for the synthesis of organic substances in plants (Renger, 2007; von Caemmerer & Baker, 2006). Increasing the nutrient absorption rate can provide the plant with sufficient nutrients for the synthesis of various physiologically active substances to cope with drought stress (Ayyaz et al., 2024). Therefore, improving the root absorption capacity can significantly improve the drought resistance of cotton. These physiological processes directly or indirectly affect each other, forming a complex network structure that jointly improves the drought resistance of cotton.

## Conclusion

This study provides compelling evidence for the role of melatonin in enhancing drought tolerance in cotton, specifically through its modulation of root cortical activity. Drought stress significantly reduces the active root cortical area, impairing root functions. Notably, the number of root cortical cells is strongly correlated with root physiological activity, as well as the growth and development of both root and shoot tissues. These results underscore the potential of exogenous melatonin to mitigate drought-induced damage by increasing the active root cortical area, promoting the accumulation of osmotic regulators, and enhancing respiratory metabolism. Such improvements facilitate more efficient water and nutrient uptake, thereby enabling plants to better adapt to water-limited conditions, mitigating the negative impact of drought stress on plant growth, and ultimately improving drought resistance (Fig. 10). Therefore, increasing the area of root cortical cells through exogenous melatonin under drought stress may be a sustainable option to improve the ability of crops to access deep-layer resources and effectively alleviate the damage caused by drought to crops.

**Figure 10.**
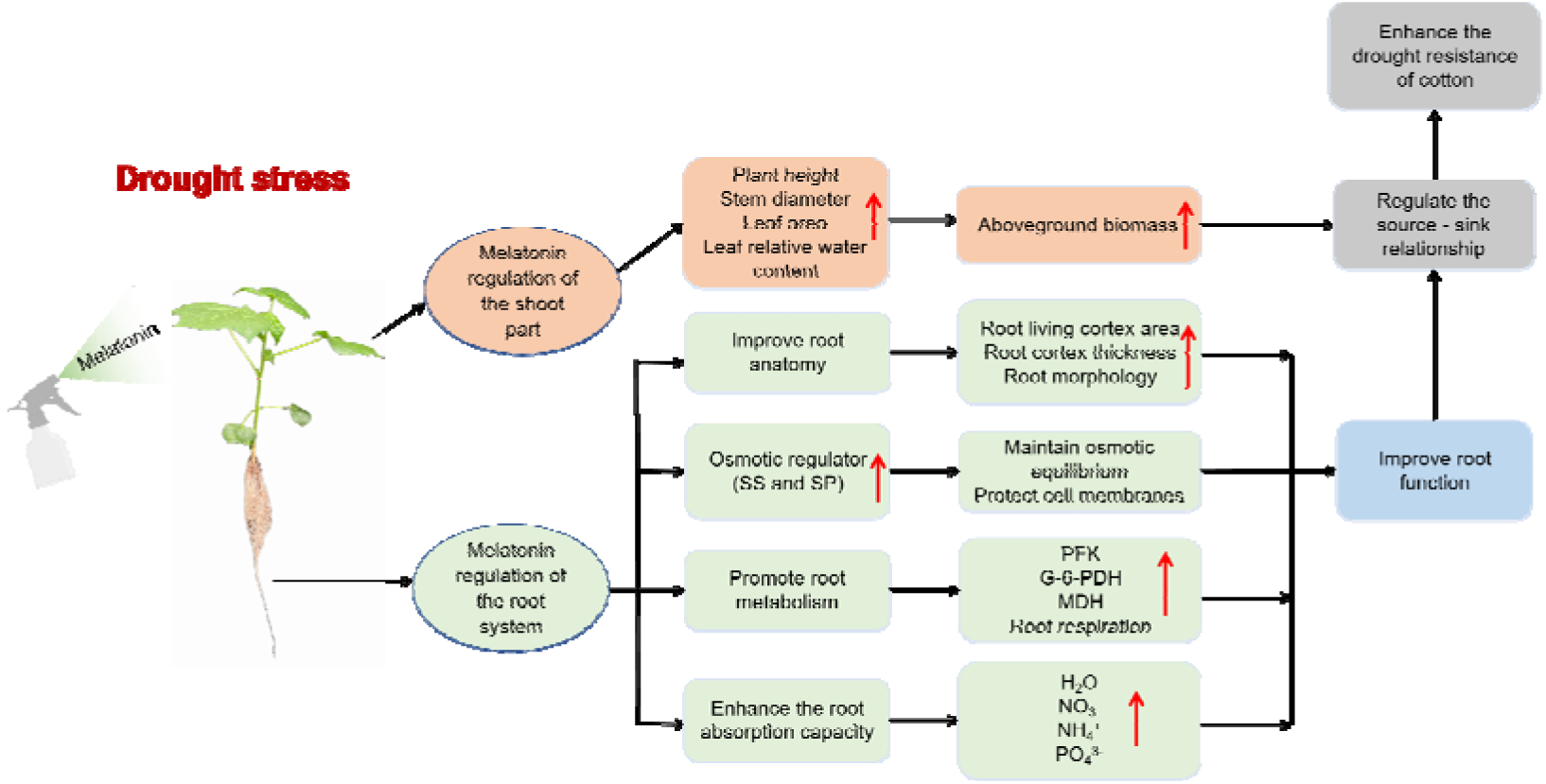
Schematic diagram of the mechanism by which exogenous melatonin enhances the drought resistance of cotton under drought stress. Note: SS: Soluble sugar content, SP: Soluble protein content, PFK: Phosphofructokinase, G-6-PDH: Glucose-6-phosphate dehydrogenase, MDH: Malate dehydrogenas

## Author Contributions

HZ and CG: Data curation, Formal analysis, Writing – original draft; LL and CL: Funding acquisition, Conceptualization, Writing – review & editing; HS, LZ, KZ, YZ, and ZW: Supervision, Investigation, Software, Resources.

## References

Ayyaz A, Batool I, Zhang K, Hannan F, Sun Y, Qin T, Athar H-R, Zafar ZU, Farooq MA, Zhou W. 2024. Unravelling mechanisms of CaO nanoparticle-induced drought tolerance in Brassica napus: an analysis of metabolite and nutrient profiling. Environmental Science: Nano 11: 2550–2567

Chen Y, Dong H. 2016. Mechanisms and regulation of senescence and maturity performance in cotton. Field Crops Research 189: 1–9

Chimungu JG, Brown KM, Lynch JP. 2014. Reduced Root Cortical Cell File Number Improves Drought Tolerance in Maize. Plant Physiology 166: 1943–1955

Cuneo IF, Knipfer T, Brodersen CR, McElrone AJ. 2016. Mechanical Failure of Fine Root Cortical Cells Initiates Plant Hydraulic Decline during Drought. Plant Physiology 172: 1669–1678

David OA, Labulo AH, Adejayan MT, Adeleke EA, Adeniyi IM, Terna AD. 2023. Anatomical adaptation of water-stressed Eugenia uniflora using green synthesized silver nanoparticles and melatonin. Microscopy Research and Technique 86: 648–658

Dong H, Niu Y, Li W, Zhang D. 2008. Effects of cotton rootstock on endogenous cytokinins and abscisic acid in xylem sap and leaves in relation to leaf senescence. Journal of Experimental Botany 59: 1295–1304

Du P, Lynch J, Sun Z, Li F-M. 2024. Does root respiration and root anatomical traits affect crop yield under stress? A meta-analysis and experimental study. Plant and Soil 1–15

Duan W, Lu B, Liu L, et al. 2022. Effects of Exogenous Melatonin on Root Physiology, Transcriptome and Metabolome of Cotton Seedlings under Salt Stress. International Journal of Molecular Sciences 23: 9456

Dubbels R, Reiter RJ, Klenke E, Goebel A, Schnakenberg E, Ehlers C, Schiwara HW, Schloot W. 1995. Melatonin in edible plants identified by radioimmunoassay and by high performance liquid chromatography-mass spectrometry. Journal of Pineal Research 18: 28–31

Dwivedi AK, Singh V, Anwar K, Pareek A, Jain M. 2023. Integrated transcriptome, proteome and metabolome analyses revealed secondary metabolites and auxiliary carbohydrate metabolism augmenting drought tolerance in rice. Plant Physiology and Biochemistry 201: 107849

Fathi A, Barari D. 2016. Effect of Drought Stress and its Mechanism in Plants. International Journal of Life Sciences 10: 1

Griffiths M, Roy S, Guo H, Seethepalli A, Huhman D, Ge Y, Sharp RE, Fritschi FB, York LM. 2021. A multiple ion-uptake phenotyping platform reveals shared mechanisms affecting nutrient uptake by roots. Plant Physiology 185: 781–795

Guo Y, Huang G, Guo Q, Peng C, Liu Y, Zhang M, Li Z, Zhou Y, Duan L. 2023. Increase in root density induced by coronatine improves maize drought resistance in North China. The Crop Journal 11: 278–290

Henry C, Deacon J. 1981. Natural (non-pathogenic) death of the cortex of wheat and barley seminal roots, as evidenced by nuclear staining with Acridine orange. Plant and Soil 60: 255–274

Hu B, Henry A, Brown KM, Lynch JP. 2014. Root cortical aerenchyma inhibits radial nutrient transport in maize (Zea mays). Annals of Botany 113: 181–189

Jaramillo RE, Nord EA, Chimungu JG, Brown KM, Lynch JP. 2013. Root cortical burden influences drought tolerance in maize. Annals of Botany 112: 429–437

Lambers H, Shane MW, Cramer MD, Pearse SJ, Veneklaas EJ. 2006. Root Structure and Functioning for Efficient Acquisition of Phosphorus: Matching Morphological and Physiological Traits. Annals of Botany 98: 693–713

Liang B, Ma C, Zhang Z, Wei Z, Gao T, Zhao Q, Ma F, Li C. 2018. Long-term exogenous application of melatonin improves nutrient uptake fluxes in apple plants under moderate drought stress. Environmental and Experimental Botany 155: 650–661

Liljeroth E. 1995. Comparisons of early root cortical senescence between barley cultivars, *Triticum* species and other cereals. New Phytologist 130: 495–501

Liu L, Guo C, Zhang K, Sun H, Zhu L, Zhang Y, Wang G, Li A, Bai Z, Li C. 2024. Root cortical senescence enhances drought tolerance in cotton. Plant, Cell & Environment 48: 615–633

Lobell DB, Roberts MJ, Schlenker W, Braun N, Little BB, Rejesus RM, Hammer GL. 2014. Greater Sensitivity to Drought Accompanies Maize Yield Increase in the U.S. Midwest. Science 344: 516–519

Lynch JP. 2015. Root phenes that reduce the metabolic costs of soil exploration: opportunities for 21st century agriculture. Plant, Cell & Environment 38: 1775–1784

Ma Z, He S, Wang X, et al. 2018. Resequencing a core collection of upland cotton identifies genomic variation and loci influencing fiber quality and yield. Nature Genetics 50: 803–813

Mukherjee S, David A, Yadav S, Baluška F, Bhatla SC. 2014. Salt stress-induced seedling growth inhibition coincides with differential distribution of serotonin and melatonin in sunflower seedling roots and cotyledons. Physiologia Plantarum 152: 714–728

Ozturk M, Turkyilmaz Unal B, García-Caparrós P, Khursheed A, Gul A, Hasanuzzaman M. 2021. Osmoregulation and its actions during the drought stress in plants. Physiologia Plantarum 172: 1321–1335

Qiao Y, Ren J, Yin L, Liu Y, Deng X, Liu P, Wang S. 2020. Exogenous melatonin alleviates PEG-induced short-term water deficiency in maize by increasing hydraulic conductance. BMC Plant Biology 20: 218

Renger G. 2007. Oxidative photosynthetic water splitting: Energetics, kinetics and mechanism. Photosynthesis Research 92: 407–25

Roy S, Griffiths M, Torres-Jerez I, et al. 2021. Application of Synthetic Peptide CEP1 Increases Nutrient Uptake Rates Along Plant Roots. Frontiers in Plant Science 12: 793145

Schneider HM, Lynch JP. 2018. Functional implications of root cortical senescence for soil resource capture. Plant and Soil 423: 13–26

Schneider HM, Postma JA, Wojciechowski T, Kuppe C, Lynch JP. 2017. Root Cortical Senescence Improves Growth under Suboptimal Availability of N, P, and K. Plant Physiology 174: 2333–2347

Sidhu JS, Lopez-Valdivia I, Strock CF, Schneider HM, Lynch JP. 2024. Cortical parenchyma wall width regulates root metabolic cost and maize performance under suboptimal water availability. Journal of experimental botany 75: 5750–5767

Singh TN, Aspinall D, Paleg LG. 1972. Proline accumulation and varietal adaptability to drought in barley: a potential metabolic measure of drought resistance. Nature New Biology 236: 188–190

Sun C, Liu L, Wang L, Li B, Jin C, Lin X. 2021. Melatonin: A master regulator of plant development and stress responses. Journal of Integrative Plant Biology 63: 126–145

Sun C, Sun N, Ou Y, Gong B, Jin C, Shi Q, Lin X. 2022. Phytomelatonin and plant mineral nutrition. Journal of Experimental Botany 73: 5903–5917

Tanaka N, Kato M, Tomioka R, Kurata R, Fukao Y, Aoyama T, Maeshima M. 2014. Characteristics of a root hair-less line of Arabidopsis thaliana under physiological stresses. Journal of Experimental Botany 65: 1497–1512

Tian J, Sethi A, Swanson BI, Goldstein B, Gnanakaran S. 2013. Taste of sugar at the membrane: thermodynamics and kinetics of the interaction of a disaccharide with lipid bilayers. Biophysical Journal 104: 622–632

Turk H, Genisel M. 2020. Melatonin-related mitochondrial respiration responses are associated with growth promotion and cold tolerance in plants. Cryobiology 92: 76–85

von Caemmerer S, Baker N. 2006. The Biology of Transpiration. From Guard Cells to Globe. Plant physiology. 143: 3–3

Wang J, Chen J, Sharma A, Tao S, Zheng B, Landi M, Yuan H, Yan D. 2019. Melatonin Stimulates Activities and Expression Level of Antioxidant Enzymes and Preserves Functionality of Photosynthetic Apparatus in Hickory Plants (Carya cathayensis Sarg.) under PEG-Promoted Drought. Agronomy 9: 702

Wang K, Xing Q, Ahammed GJ, Zhou J. 2022. Functions and prospects of melatonin in plant growth, yield, and quality. Journal of Experimental Botany 73: 5928–5946

Xiao P-X, Chen X, Zhong N-Y, Zheng T, Wang Y-M, Wu G, Zhang H, He B. 2023. Response of Vicia faba to short-term uranium exposure: chelating and antioxidant system changes in roots. Journal of Plant Research 136: 413–421

Yancey PH. 2005. Organic osmolytes as compatible, metabolic and counteracting cytoprotectants in high osmolarity and other stresses. The Journal of Experimental Biology 208: 2819–2830

Yang K, Sun H, Liu M, et al. 2023. Morphological and Physiological Mechanisms of Melatonin on Delaying Drought-Induced Leaf Senescence in Cotton. International Journal of Molecular Sciences 24: 7269

Yang Z, Tian J, Wang Z, Feng K. 2022. Monitoring the photosynthetic performance of grape leaves using a hyperspectral-based machine learning model. European Journal of Agronomy 140: 126589

Zhan A, Lynch JP. 2015. Reduced frequency of lateral root branching improves N capture from low-N soils in maize. Journal of Experimental Botany 66: 2055–2065

Zhou K, Cheng Q, Dai J, Liu Y, Liu Q, Li R, Wang J, Hu R, Lin L. 2023. Effects of exogenous melatonin on sugar and organic acid metabolism in early-ripening peach fruits. PLOS ONE 18: e0292959

Zhu J, Schwörer S, Berisa M, Kyung YJ, Ryu KW, Yi J, Jiang X, Cross JR, Thompson CB. 2021. Mitochondrial NADP(H) generation is essential for proline biosynthesis. Science 372: 968–972

Zhu L, Li A, Sun H, et al. 2023. The effect of exogenous melatonin on root growth and lifespan and seed cotton yield under drought stress. Industrial Crops and Products 204: 117344

Zhu L, Sun H, Wang R, et al. 2024. Exogenous melatonin improves cotton yield under drought stress by enhancing root development and reducing root damage. Journal of Integrative Agriculture 23: 3387–3405

